# Propylene glycol inactivates respiratory viruses and prevents airborne transmission

**DOI:** 10.1101/2023.02.13.528349

**Authors:** Christine T. Styles, Jie Zhou, Katie E. Flight, Jonathan C. Brown, Michael Vanden Oever, Thomas P. Peacock, Ziyin Wang, Rosie Millns, John S. O’Neill, Wendy S. Barclay, John S. Tregoning, Rachel S. Edgar

## Abstract

Viruses are vulnerable as they transmit between hosts and we aimed to exploit this critical window. We found that the ubiquitous, safe, inexpensive and biodegradable small molecule propylene glycol (PG) has robust virucidal activity. Propylene glycol rapidly inactivates influenza, SARS-CoV-2 and a broad range of other enveloped viruses, and reduces disease burden in mice when administered intranasally at concentrations commonly found in nasal sprays. Most critically, aerosolized PG efficiently abolishes influenza and SARS-CoV-2 infectivity within airborne droplets, potently preventing infection at levels significantly below those well-tolerated by mammals. We present PG vapor as a first-in-class non-toxic airborne virucide, to prevent transmission of existing and emergent viral pathogens, with clear and immediate implications for public health.

**One-sentence summary:** Propylene glycol is a potent and safe virucidal compound that could be used to limit and control infections.

## Main Text

The COVID-19 pandemic has claimed >6.6 million lives so far*(1)*, and the World Health Organization estimates seasonal influenza mortality at 290,000 – 650,000 people annually *(2)*. Beyond this health burden, respiratory viruses cause severe economic and societal costs, recently estimated for the UK government alone at £23 billion/year during future influenza-type pandemics and £8 billion/year for seasonal influenza *(3)*. Public health and social measures used to combat respiratory virus transmission include mask wearing, physical distancing, lockdown and travel restrictions *(4)*. Such measures are primarily evidenced by observational studies rather than randomised controlled trials*(5, 6),* and require compliance*(7, 8).* Other strategies involve improving ventilation and frequent disinfection to remove infectious virus from the environment, but both come with significant drawbacks and there is growing concern over the health and environmental consequences of prolonged, widespread use of disinfectants*(9–13)*. Natural ventilation is not always suitable, increasing risk from air pollutants and vector-borne diseases along with thermal and energy considerations in colder climates. Expensive engineering solutions like mechanical ventilation require co-ordinated action across government, health, transport, business, housing and environmental sectors, and a significant culture shift to prioritise infection resilience in indoor environments alongside net zero objectives*(3)*. There is an urgent unmet need for novel nonpharmaceutical interventions to combat the spread of emerging pathogens and seasonal diseases.

Propylene glycol (PG, propan-1,2-diol) is a synthetic liquid compound, whose amphiphilic properties are utilised in a wide range of products and industries: food and drink, cosmetics and pharmaceuticals, including oral, topical, intravenous and nebulised drug delivery*(14)*. PG is considered a ‘<u>G</u>enerally <u>R</u>ecognised <u>A</u>s <u>S</u>afe’ (GRAS) molecule, efficiently metabolised and excreted from mammals, and is approved for widespread applications by the US Food and Drug Agency (FDA) and European Medicines Agency (EMA)*(14, 15).* PG is largely used as a vehicle or humectant in these preparations due to its water-absorbing properties, however it has both anti-bacterial*(16–18)* and anti-fungal activity*(19),(20)*. Studies conducted in the 1940s show PG vapor reduced the infectivity of aerosolized bacterial pathogens in mouse models, preventing sepsis-induced mortality*(16, 17, 21)*. In human trials from the same era, vaporized PG reduced the airborne bacterial burden in army barracks*(22)*, and the introduction of PG vapor on a children’s convalescent ward reduced the incidence of undefined respiratory illnesses between 1941-44*(23).* One study suggested PG vapor could protect mice against disease when exposed to airborne influenza*(24)*, and reduce vaccinia- and influenza-mediated fatality in chick embryos*(25, 26).* These findings preceded the advent of molecular virology, however, so the impact of PG on virus particle infectivity was never directly assessed and remains poorly defined.

We tested the hypothesis that PG is virucidal and could reduce respiratory virus transmission by droplet, aerosol and fomite routes.

### PG inactivates influenza A virus and reduces disease severity

To examine whether PG exhibits virucidal activity, we first focused on influenza A virus (IAV), a causative agent of seasonal outbreaks in humans and birds that significantly burden healthcare systems and the poultry farming industry worldwide*(27)*, and responsible for the worst pandemics of the previous century*(28)*. IAV was incubated with different concentrations of PG, and infectivity was determined by titration.

PG dramatically reduced IAV infection of cultured cells, with virucidal activity dependent on both PG concentration and incubation time (Fig. 1A-C). PG-mediated IAV inactivation was temperature-dependent, with progressively greater virucidal activity evident at 32°C (nasal and skin temperature) and 37°C (body temperature) compared with 20°C (room temperature, RT). At physiological temperatures, 60% PG reduced IAV infectivity roughly 10,000-fold within 5 minutes, down to undetectable levels after 30 minutes. This demonstrates PG has potent virucidal activity.

**Fig. 1.**
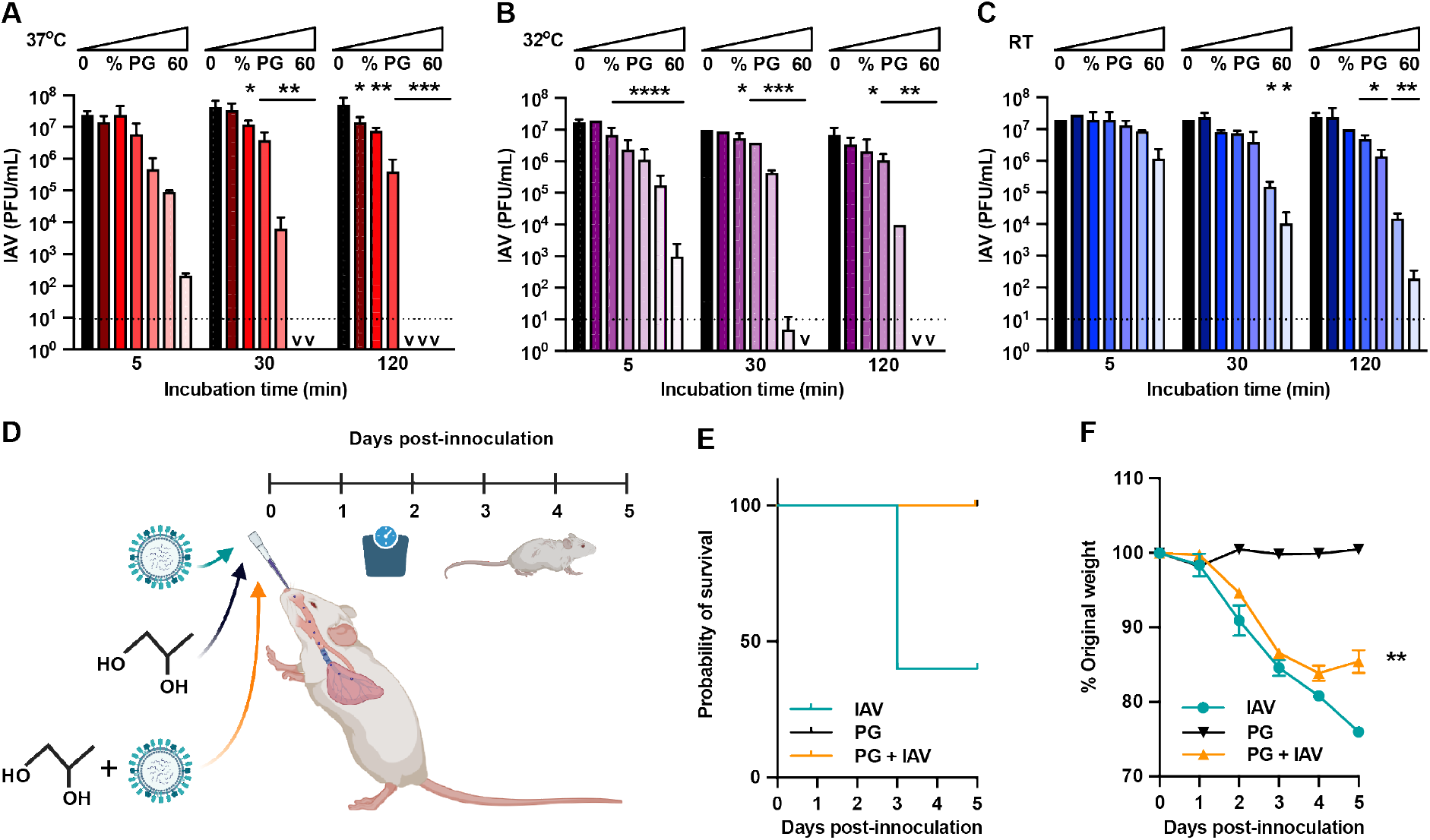
Propylene glycol (PG) reduces influenza A virus infectivity *in vitro* and *in vivo.* IAV was incubated with 10-60% PG, or medium-only control for 5-120 minutes at 37°C (**A**), 32°C (**B**) or room temperature (RT) (**C**) and infectivity assessed by plaque assay (N=2; mean±SD); PFU = plaque forming units. 2-way ANOVA ([PG] x time): (**A**) [PG] ****P<0.0001, time P>0.05, interaction P>0.05; (**B**) [PG] ****P<0.0001, time ****P<0.0001, interaction ***P<0.001; (**C**) [PG] ****P<0.0001, time *P<0.05, interaction P>0.05; multiple comparisons versus control. v = <10^1^; dashed line = limit of detection. For all multiple comparisons *P<0.05, **P<0.01, ***P<0.001, ****P<0.0001. (**D**) *In vivo* methodology (BioRender). Mice were intranasally inoculated with 50mL total volume of PG only (20% PG in PBS), H1N1 IAV only (5×10^4^ PFU in PBS), or PG+IAV and monitored for 5 days (N=5 mice/group; mean±SD). (**E**) Mouse survival after infection. (**F**) Weight loss after infection. Mixed-effect analysis: [Time] ****P<0.0001, [PG] ****P<0.0001; IAV alone *versus* PG+IAV on day 5 **P=0.0056.

To determine the translational potential of PG-mediated virucidal activity against IAV, we then investigated combined inhalation of IAV and PG *in vivo.* 20% PG was the lowest concentration to yield statistically significant reduction in IAV infectivity at nasal temperature (Fig. 1B). Therefore, mice were intranasally inoculated with 20% PG alone, IAV alone or IAV and 20% PG in combination, with disease progression tracked over 5 days (Fig. 1D). Following inhalation of IAV alongside PG, mice showed enhanced survival and reduced clinical signs compared to mice infected with IAV alone (Fig. 1E, Fig. S1). 3/5 mice within the IAV only group showed such poor clinical scores that they were humanely culled 3 days after infection. Mice also lost significantly less weight when PG was co-administered with IAV than with IAV alone (Fig. 1F), whereas mice receiving PG only showed no change in clinical scores or weight over 5 days, consistent with its long-established biological safety in mammals. No mice in the IAV+PG or PG-only group met the severity limit necessitating humane killing, demonstrating the protective nature of intranasally administered PG. We conclude that PG can safely reduce the infectivity of influenza A virus *in vitro* and *in vivo.*

### PG has broad-spectrum virucidal activity

We next asked whether PG could inactivate other enveloped viruses, including the virus responsible for the COVID-19 pandemic; severe acute respiratory syndrome coronavirus 2 (SARS-CoV-2). We found that PG inactivated the IC19 strain of SARS-CoV-2*(29)* with even greater efficiency than observed for IAV (Fig. 1A, 2A). After 1 minute treatment with 50% PG at room temperature, SARS-CoV-2 infectivity decreased by >1000-fold, indicating clear virucidal activity that persisted over longer time frames (Fig. 2A). PG also efficiently reduced the infectivity of the enveloped double stranded DNA g-herpesvirus Epstein Barr (EBV), a lifelong infection carried by most of the human population and associated with numerous cancers*(30)*. Similar to IAV and SARS-CoV-2, PG showed robust virucidal activity against EBV, with >1000-fold reduction in viral titre upon incubation with 50% PG (Fig. 2B).

**Fig. 2.**
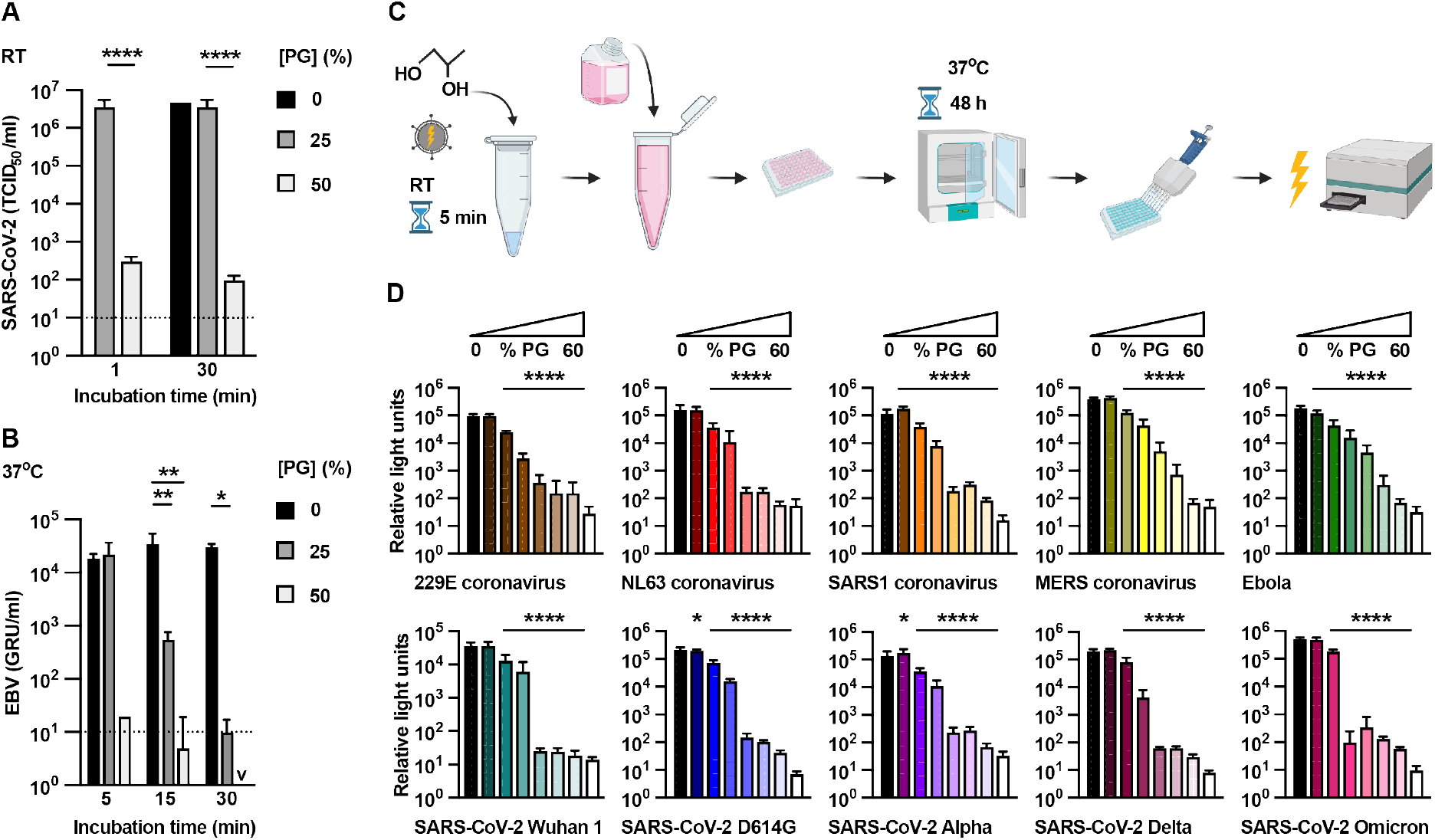
PG inactivates SARS-CoV-2, EBV and many different pseudoviruses. (**A**) SARS-CoV-2 incubated with 0-50% [PG] for 1-30min at RT and infectivity assessed by TCID50 assay (N=2, n=4; mean±SD). 2-way ANOVA ([PG] x time): [PG] *P<0.05, time P>0.05, interaction P>0.05. Dotted line = limit of detection. (**B**) EBV incubated with 0-50% [PG] for 5-30min at 37°C and infectivity assessed by titration (N=2; mean±SD; GRU=green Raji units). 2-way ANOVA ([PG] x time): [PG] ****P<0.0001, time P>0.05, interaction P>0.05; v = <10^1^. Methodology (BioRender) (**C**) and results (**D**) of lentivirus pseudotypes expressing different viral glycoproteins incubated with 0-60% PG for 5min at RT before assessing infectivity by bioluminescence (N=2, n=3; mean±SD). White bars = mock infected controls. 1-way ANOVA [PG] compared to 0% PG control. For all multiple comparisons *P<0.05, **P<0.01, ***P<0.001, ****P<0.0001.

To explore the broader context of PG’s activity against disease-causing viruses, we employed a pseudovirus system engineered to express viral envelope glycoproteins from diverse human pathogens, including NL63 and 229E seasonal coronaviruses, severe acute respiratory syndrome coronavirus (SARS), middle eastern respiratory syndrome coronavirus (MERS) and Ebola. Using this platform, we also tested PG against glycoproteins from different SARS-CoV-2 variants, including the most recent variant of concern: Omicron.

Using a bioluminescence-based assay for infectivity (Fig. 2C), PG significantly limited the infection capability of every different pseudovirus, rapidly reducing entry into susceptible cells in a dose-dependent manner (Fig. 2D, Fig. S2). Although PG consistently reduced infectivity, the concentration required varied between the different pseudovirus-expressed glycoproteins: recapitulating the variation in the specific potency of PG against IAV, SARS-CoV-2 and EBV virus particles. This strongly suggests different PG virucidal thresholds and modes of action against specific viruses, against an overall background of broad-spectrum virucidal activity against enveloped viruses.

### Vaporized PG inactivates airborne viruses

Respiratory droplets and aerosols that are exhaled/expelled by talking, sneezing or coughing from infected individuals represent a major transmission route for many pathogens, particularly respiratory viruses such as influenza and SARS-CoV-2*(31)*. Artificially generated aerosols of SARS-CoV-2 and IAV remain infectious for at least 3h and 1h, respectively*(32),(33)*, with viable SARS-CoV-2 aerosols identified at >2m distance from infectious patients*(34)*. The COVID-19 pandemic has highlighted the clear and pressing need for effective, safe and economical ways to inactivate infectious particles from contaminated air. Current virucidal disinfectants are unsafe for human consumption and often environmentally harmful*(9–13)*. PG is biodegradable and non-toxic, with numerous studies showing PG vapor can be safely inhaled for long durations without adverse effects, testing up to 41 mg PG/L air *(35–41).*

The condensation of vaporized or aerosolised PG with airborne aqueous respiratory droplets is very energetically favourable, and occurs rapidly in atmospheric air at room temperature. We predicted that low levels of vaporized PG would condense with respiratory droplets in sufficient amounts to inactivate any airborne virus particles therein. To model infection by airborne IAV and SARS-CoV-2 we used a bespoke transmission tunnel system (Fig. 3A)*(42)*. Within the transmission tunnel, permissive cell monolayers at different distances were exposed to airborne virus droplets (4-6μm) in the presence of total vaporized PG concentrations from 0 - 11mg/L air (Fig. S3). Following exposure, viral plaque area was computationally derived *via* two independent methods. In line with our predictions and the 1941 pathogenesis study*(24)*, PG vapor reduced airborne IAV and SARS-CoV-2 infectivity in a dose-dependent manner (Fig. 3B-F, Fig. S4), abolishing infection within a distance of <1m. Vapor was a more efficacious virucide than PG in solution (Fig. 1A-C, Fig. 2), as was PG within nebulised droplets (Fig. S5A,B,C *versus* Fig. S5D), mirroring inhalation in mice (Fig. 1D-F).

**Fig. 3.**
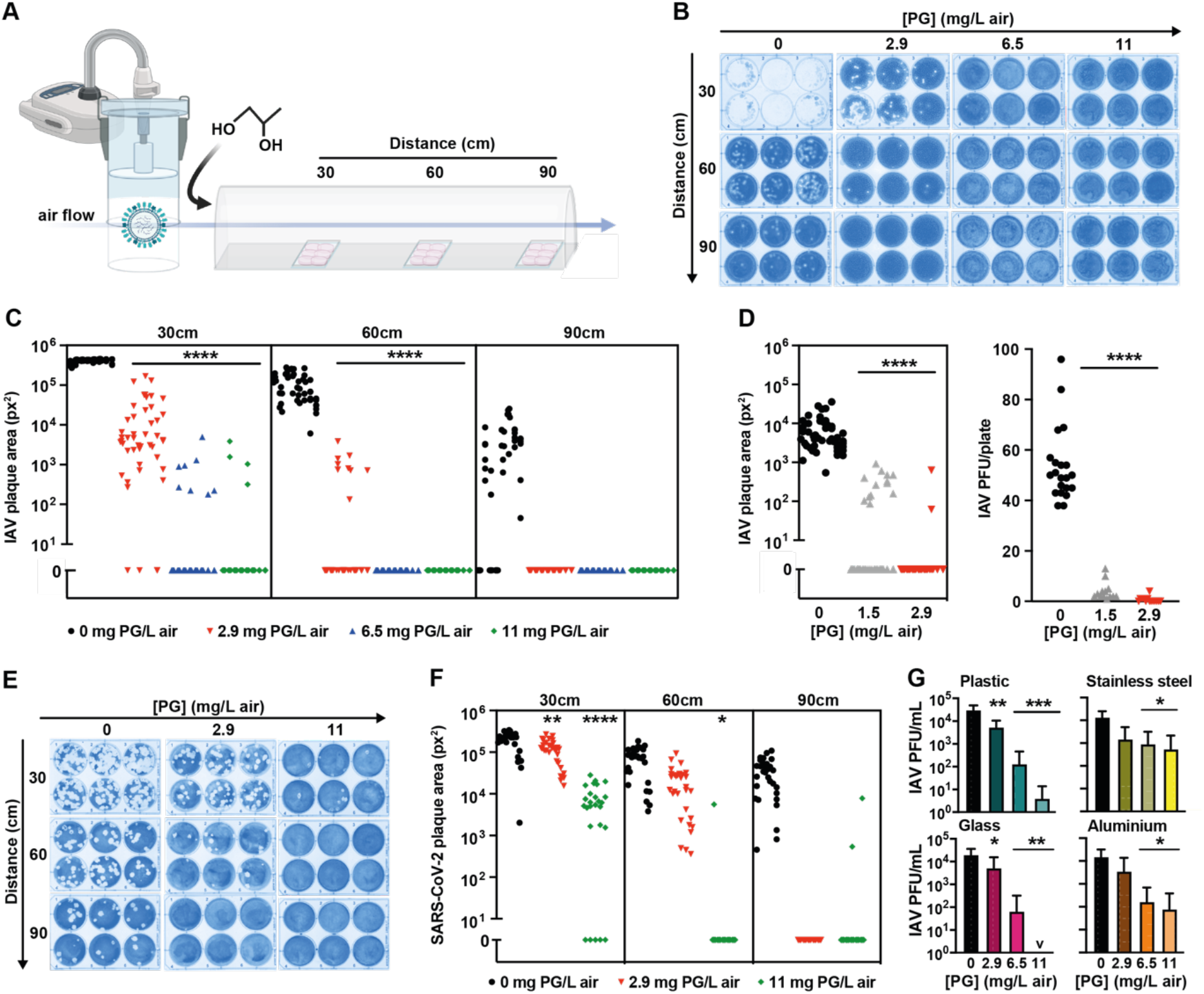
PG vapor efficiently inactivates airborne IAV and SARS-CoV-2. (**A**) Virus transmission tunnel schematic (BioRender). (**B**) PG vapor was introduced into the transmission tunnel to a final concentration of 0-11 mg/L air prior to nebulization of 10^6^ PFU IAV. Representative IAV culture plates; (**C**) viral plaque area on culture plates at 30cm, 60cm and 90cm distance was computationally analysed using ImageJ ColonyArea plugin (N=8, n=6); 2-way ANOVA ([PG] x distance): [PG] ****P<0.0001, distance ****P<0.0001, interaction **** P<0.0001. (**D**) PG vapor was introduced into the transmission tunnel to a final PG concentration of 0-2.9 mg/L air prior to nebulisation of 10^4^ PFU IAV. Viral plaque area at 30cm distance analysed as for (B)(N=8, n=6; 1-way ANOVA [PG]: ****P<0.0001) and plaques counted (1-way ANOVA [PG]: ****P<0.0001). (**E**) PG vapor was introduced into the transmission tunnel to a final concentration of 0-11 mg/L air prior to nebulization of 3×10^4^ PFU SARS-CoV-2 Delta variant. Representative SARS-CoV-2 culture plates; (**F**) viral plaque area was assessed as per (B)(N=5, n=6); 2-way ANOVA ([PG] x distance): [PG] ***P<0.001, distance ****P<0.0001, interaction *** P<0.001. (G) 10^5^ PFU IAV on fomite model surfaces (plastic, stainless steel, aluminium and glass) were exposed to vaporized PG (0-11 mg/L air) and infectivity assessed by plaque assay after 25min (N≥4, n≥3; mean±SD). 1-way ANOVA [PG]: *P<0.05, **P<0.01, ***P<0.001, ****P<0.0001.

Whilst the transmission tunnel has some limitations as a model of viral dissemination through droplets and aerosols (see Fig. S4), our findings clearly demonstrate efficient and rapid PG-mediated inactivation of airborne IAV and SARS-CoV-2, at virus levels that far exceed the estimated amounts expelled by speaking, coughing or sneezing*(42–44)*. 1.5 mg PG/L air was the lowest exposure we could consistently generate with our experimental system, and it effectively abolished infectivity when less IAV was nebulized into the tunnel to mimic an amount comparable to multiple human coughs (Fig. 3D, Fig. S4B). Given the strong correlation between initial viral dose, infection probability and disease *severity(45–50),* clinical studies are now required to identify the optimum aerosolized PG levels that effectively reduce viral transmission under ‘real-world’ conditions.

Alongside airborne routes, viruses are also transmitted indirectly *via* contact with surfaces contaminated by the deposition of virus-containing respiratory droplets and subsequent mechanical transfer to mucous membranes (fomite transmission)*(31)*. Infectious SARS-CoV-2 and SARS-CoV can be recovered up to 72 hours after deposition on surfaces, including plastic and stainless steel*(32)*, and viable H1N1 IAV is recoverable for up to 2 weeks from stainless steel*(51)*. Importantly then, PG vapor inactivated IAV upon varied fomites, with PG at 11mg/L air sufficient to significantly reduce infectious IAV on all surfaces tested (Fig. 3G). As such, we propose PG as the first-in-class example of a safe, inexpensive and environmentally neutral broad-spectrum virucide, for inactivation of airborne and surface-bound viruses.

## Discussion

With the increasing threat from emerging pathogens, we need new tools that can immediately be deployed to attenuate viral transmission within social, healthcare and transport settings. For example, sanitizing ambulances between patients caused delays and disruption during the COVID-19 pandemic*(55)*. PG vapor is a potentially valuable resource for limiting diverse virus infections by multiple routes, including aerosol, droplet and fomite transmission; an economical and effective virucide that is much safer to ingest and inhale compared to other disinfectants and fumigation systems, whilst also avoiding their negative environmental consequences and toxicity*(9)*. Prospective application of PG as an infection prevention measure requires further investigation beyond laboratory settings, but our results strongly suggest we should leverage its virucidal capacity. PG is already an excipient in nonprescription nasal sprays so this intervention could be rapidly implemented, and vapor generation utilizes existing technologies.

It was beyond the scope of this study to determine whether PG reduces infectivity of non-enveloped viruses. We predict that PG is disrupting the lipid envelope, as shown for bacterial membranes*(52)*. However, variation in the PG concentration required to prevent entry of pseudoviruses expressing different envelope glycoproteins (Fig. 2) suggests that PG may induce conformational changes that can also restrict infection. Therefore, PG may exert some virucidal effect against non-enveloped viruses by altering the structure of surface proteins that mediate host membrane penetration or by direct capsid disruption*(53)*. Most conventional antiseptics and disinfectants used within hospitals such as ethanol hand gels exhibit poor activity against non-enveloped viruses like human norovirus and rotaviruses*(54)*, so it is of commercial and pharmaceutical interest to assess whether PG can reduce their infectivity. Recently, PG was shown to disrupt rotavirus replication factories inside cells, demonstrating the potential sensitivity of phase separated viroplasms to this small molecule*(55)*. In addition, studies from the 1940s suggest PG vapor is bactericidal against many aerosolised bacterial species *(16, 17, 21)*. Re-evaluating these findings using modern experimental methodologies is paramount, given the threat from antimicrobial resistance. We think PG acts *via* biophysical disruption of lipid membranes or protein structure, presenting a substantial, if not insurmountable, barrier to evolutionary escape by pathogens.

## Conclusion

PG is already approved for use within pharmaceutical, cosmetic and food industries, yet its inherent virucidal activity has not been examined or exploited. PG in nasal sprays, nebulizers and sanitizers would protect vulnerable individuals, and direct inactivation of airborne human viruses by PG vapor could potentially reduce the overall infectious burden and transmission rates in clinical and commercial settings.

## Materials and methods

### Cell culture

All media and supplements were supplied by Gibco-Life Technologies and cells were maintained in a humidified incubator at 37°C with 5% CO2. Madin-Darby Canine Kidney (MDCK), African green monkey kidney (Vero E6), human hepatoma-derived 7 cell line (Huh7) and human embryonic kidney cells (293T) transduced with an ACE2 lentiviral vector (ACE2-293T, as described previously*(56)*) were routinely cultured in Dulbecco’s modified Eagle’s Medium (DMEM). Raji cells were cultured in RPMI-1640 medium (RPMI). Media was supplemented with 10% foetal calf serum (FCS), Glutamax and penicillin/streptomycin. ACE2-293T cells were additionally supplemented with 1 ug/mL puromycin. Vero E6 cells overexpressing ACE2 and TMPRSS2 (VAT cells, as described previously*(57)*), were additionally supplemented with non-essential amino acids, 0.2 mg/mL Hygromycin B and 2 mg/mL Geneticin^TM^ (G418 Sulfate).

### IAV infectivity plaque assay

Influenza strain A/PR8/8/34 (H1N1) was used in this study and referred to as IAV. IAV was propagated in confluent MDCK cells in the presence of 1 mg/mL TPCK treated trypsin (Worthingtons Bioscience) in serum free medium (SFM)(DMEM, penicillin/streptomycin and Glutamax). IAV was incubated with 0 – 60% v/v concentration of propylene glycol (PG)(Sigma) for 5-120 minutes as described, at either room temperature (RT) or at 32°C and 37°C. Following PG treatment, the virus/PG suspensions were serially diluted in SFM with the initial 10^-1^ dilution being the detection limit for IAV infectivity in this assay. Confluent MDCK cells were incubated with each serial dilution for 1h at 37°C then input virus was removed by aspiration and cells overlaid with SFM containing 0.14% BSA (Sigma), 0.8% Avicel© (FMC BioPolymer) and 1 mg/mL TPCK treated trypsin. After 72h, cells were fixed in 8-10% formalin/PBS, stained with 0.1% toluidine blue (Sigma) and viral plaque forming units (PFU) assessed.

### IAV fomite infectivity assay with PG vapor

2μL droplets containing 10^5^ PFU IAV in SFM were pipetted onto stainless steel discs, polystyrene plastic, aluminium foil sections or glass discs within 6 well plates. Plates were placed in a sealed polystyrene chamber and exposed to vaporized PG (0 – 11 mg/L air) using a MicroFogger 2 (WorkshopScience). After 25 min, virus was recovered in 1 mL SFM, serially diluted and quantified by plaque assay on MDCK as described above.

### EBV infectivity assay

Prototypical laboratory strain Epstein Barr Virus (EBV) containing a GFP cassette was incubated with 0 – 50% v/v PG for 5 - 120 min at 37°C. Following treatment, virus/PG suspensions were serially diluted in RPMI with the initial 10^-1^ dilution being the detection limit for EBV infectivity in this assay. 5×10^4^ Raji cells were added to each dilution and incubated for 48h at 37°C. RPMI containing 20 nM TPA and 5 mM sodium butyrate was added and a viral titre determined after 24h using fluorescent microscopy to identify GFP-expressing cells (Green Raji units; GRU).

### Lentivirus pseudotype infectivity assay

Pseudotype lentiviruses were generated in HEK 293T cells as described previously*(56)*. Briefly, 293T cells were co-transfected with plasmids encoding desired envelope glycoprotein, firefly luciferase reporter (pCSGW) and pCAGGs-GAG-POL using Lipofectamine 3000 (Thermofisher) and pseudovirus harvested at 48h and 72h post-transfection. A control pseudovirus was also constructed without a viral glycoprotein component (‘bald’ pseudovirus; Fig. S2). Pseudoviruses used in this study contained glycoproteins from 5 different SARS-CoV-2 variants (Wuhan-1*(29)*, D614G*(58),*, Alpha*(58)*, Delta*(59)* and Omicron*(59)*), middle eastern respiratory syndrome coronavirus (MERS-CoV)*(29)*, SARS-CoV*(29)*, NL63 and 229E coronaviruses*(29)*, Ebola*(60)*, amphotropic murine leukaemia virus (MLV-A)*(60)* or Indiana vesicular stomatitis virus (VZV-G)*(60)*.

Pseudoviruses were treated with 0 – 60% v/v [PG] for 5 min at room temperature then diluted in growth media. Pseudoviruses were plated in triplicate onto confluent ACE2-293T cells (SARS-CoV-2 variants, SARS-CoV, NL63, Ebola, MLV-A, VZV-G and ‘bald’) or Huh7 cells (MERS-CoV and 229E) and incubated for 48h. Luciferase activity was measured using a Firefly luciferase assay system kit (Promega), on a FLUOstar Omega plate reader (BMF labtech). Each analysed plate contained triplicate uninfected cells to control for background luminescence.

### SARS-CoV-2 TCID50 infectivity assay

The strain of SARS-CoV-2 used for infectivity assays was SARS-CoV-2/England/IC19 and is henceforth referred to as ‘SARS-CoV-2’ *(29).* Vero E6 cells in assay diluent (DMEM, 0.3% BSA, NEAA, penicillin/streptomycin) were seeded into 96-well plates and incubated at 37°C for 24h. SARS-CoV-2 was incubated with 0 - 50% v/v PG for 1 – 30 min at room temperature. Virus/PG suspension was then added to the first column of confluent Vero E6 cells and a log_10_/half-log_10_ dilution series immediately performed in assay diluent. Technical replicates were performed for each sample. Plates were incubated for 5 days before adding an equal volume of crystal violet stain (0.1% w/v) to live cells. Wells were scored for either an intact, stained cell sheet or the absence of cells due to virus-induced cytopathic effect. For each condition, the Spearman-Karber method was used to calculate the 50% tissue culture infectious dose (TCID50) of virus.

### Virus transmission tunnel experiments with airborne IAV and SARS-CoV-2

A custom-built transmission tunnel system described by Singanayagam *et al(42)*(Fig. 3A) was used to assess aerosolized PG-mediated inactivation of nebulised IAV and SARS-CoV-2 (B.1.617.2, Delta variant*(61)*) viruses. Briefly, the transmission tunnel holds 3 tissue culture plates at intervals of 30cm, 60cm and 90cm from the nebulizer (Aerogen Pro nebuliser; Aerogen). A bias flow pump maintains directional air flow from the nebulizer chamber to the exposure tunnel that deposits airborne virus within 4-6μm diameter droplets across the tissue culture plates during a 30min exposure period.

All transmission tunnel experiments were performed within a class I (SARS-CoV-2) or class II (IAV) biological safety cabinet. MDCK or VAT cells were seeded into 6 well plates for IAV or SARS-CoV-2 analysis, respectively. Before placing in the transmission tunnel, MDCK cells were transferred to SFM and VAT cells were transferred to assay medium (MEM, penicillin/streptomycin, 4 mM L-glutamine, 0.4% (w/v) BSA, 0.32% NaHCO2 and 20 mM HEPES). To assess inactivation of IAV and SARS-CoV-2 by aerosolized PG, concentrations between 0 - 11 mg/L air were introduced into the exposure tunnel using a MicroFogger 2 (WorkshopScience). To quantify the total airborne PG concentration (vapor plus droplets), sampling was performed using a SKC biosampler attached to the tunnel that collected PG into 1 mL distilled water. Samples were analysed on an Osmomat 3000 (Gonotec) and total PG concentration calculated by comparison to a standard curve of PG concentrations (Fig. S3). After PG was aerosolized within the tunnel, permissive cells were exposed to either 10^4^ - 10^6^ PFU nebulized IAV in PBS or 3 x 10^4^ nebulized PFU SARS-CoV-2 in PBS by starting the directional air flow for 30 min. Cells were then incubated for 1h at 37°C and 5% CO2 in a humidified incubator. For IAV, exposed medium was aspirated and cells overlaid with SFM supplemented with 0.8% v/w Avicel© and before fixation and staining 72h later as previously described. For SARS-CoV-2, assay medium supplemented with Avicel© (0.75% w/v final concentration) was added directly to cells and incubated for 72h before cells were fixed and stained with crystal violet (0.1% w/v) containing 30% EtOH for >30 min.

Transmission tunnel tissue culture plates were imaged using a BioRad Gel Visualiser and each well within the plate was independently computationally analysed. Virus mediated clearance of cells (plaque area) was quantified using two independent ImageJ FIJI platform systems; the plugin ColonyArea*(62)* and the macro ViralPlaque*(63)*. ColonyArea quantifies the % well area containing stained cells and ViralPlaque quantifies the % area lacking stained cells due to viral cytopathic effect. Calculations were consistently between the two methods (Fig. 3 and Fig. S4; Fig. S4D for correlation). Data is presented as plaque area in square pixels (px^2^).

To assess PG-mediated inactivation of IAV during aerosolization (Fig. S5), 0% or 40% PG was added to 10^6^ PFU then immediately nebulised into the chamber (<1 min incubation at room temperature), MDCK cells exposed to the aerosolised PG/IAV mixture and infectious IAV quantified as above.

### Combined inhalation of PG and IAV *in vivo*

6-8 week old female BALB/c mice were obtained from Charles River UK Ltd (Portscatho, UK) and kept in specific-pathogen-free (SPF) conditions in accordance United Kingdom’s Home Office guidelines. All work was approved by the Animal Welfare and Ethical Review Board (AWERB) at Imperial College London. For infections, mice were anesthetized using isoflurane and inoculated intranasally (i.n.) with 50 μL final volume of either 5×10^4^ PFU H1N1 A/California/7/2009 influenza virus (IAV), 5×10^4^ IAV and 20% PG, or 20% PG alone in sterile PBS. Mice were weighed after infection. Mice were scored for signs of clinical illness after infection based on an adapted scoring system*(64)*. Scores were given (0 to 5, with 0 being in healthy condition) for coat, activity, stance and breathing. Mice were culled using 100 μl intraperitoneal pentobarbitone (20 mg dose, Pentoject, Animalcare Ltd. UK), tissues collected and cells in bronchoalveolar lavage (BAL) were counted as previously described*(65)*. Nasal lavage and BAL were processed for viral load by plaque assay on MDCK cells as described above.

### Statistical analysis

Data was analysed in GraphPad Prism, and presented as mean±standard deviation (SD). At least 2 independent biological replicates were performed for each experimental condition. Detailed information for one way- and two way-ANOVA analyses including F values, degrees of freedom and replicate details are shown in Table S1. Statistical test outcomes including multiple comparisons are summarized in Table S2.

## Acknowledgments

The authors thank R Frise and A Borodavka for discussion, R White and P Farrell for Raji cells and P McKay for codon-optimized spike proteins used to generate pseudoviruses. We also thank N Gunawardana, T Santarius, F Harris and their teams at Addenbrooke’s Hospital, Cambridge, whose care of RSE enabled this research to continue.

## Author contributions

Conceptualization: RSE

Methodology: CTS, JZ, JB, TP, WSB, JST, RSE

Investigation: CTS, JZ, KEF, JB, MVO, TP, ZW, RM

Formal Analysis: CTS, JB, JST, RSE

Visualization: CTS, RSE

Funding acquisition: JSO, WSB, JST, RSE

Project administration: JSO, WSB, JST, RSE

Supervision: CTS, JSO, WSB, JST, RSE

Writing – original draft: CTS, RSE

Writing – review & editing: CTS, JZ, KEF, JB, MVO, TP, ZW, RM, JSO, WSB, JST, RSE

## Data and materials availability

The data generated in this study is available within the article and supplementary materials. Raw image files and materials are available from the corresponding author upon request. All quantifications and statistical analyses will be made available upon publication.

## List of supplementary materials

Fig. S1

Fig. S2

Fig. S3

Fig. S4

Fig. S5

Table S1

Table S2

