## Supplementary figures 1 - 5 and tables 1 - 2 for "Propylene glycol inactivates respiratory viruses and prevents airborne transmission"

**This PDF file includes:**

Figures S1 to S5
Tables S1 and S2

**Figure S1. Concomitant inhalation of IAV and PG reduces clinical burden.**

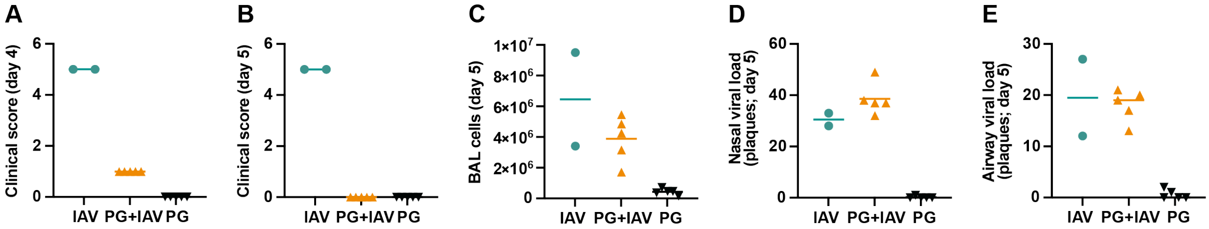


Mice were intranasally inoculated with PG alone (20% PG in PBS), H1N1 IAV alone ($5x10^4$ PFU in PBS), or PG+IAV (50µL total volume for all groups; N=5 mice/group) and monitored for 5 days (see **Fig. 1D**). Clinical scores on day 4 (**A**) and day 5 (**B**), BAL cell count (**C**), viral nasal load (**D**) and airway viral load (**E**). 3/5 mice in the IAV only group were culled prior to day 4/5 collection point due to poor clinical scores, limiting the power of this study to detect statistically significant differences between groups from day 5 *post-mortem* immunological and virological assays, but decreased clinical score and inflammatory BAL cell counts were observed in the PG+IAV group compared to IAV alone.

**Figure S2. PG exhibits virucidal activity against pseudoviruses expressing viral glycoproteins.**

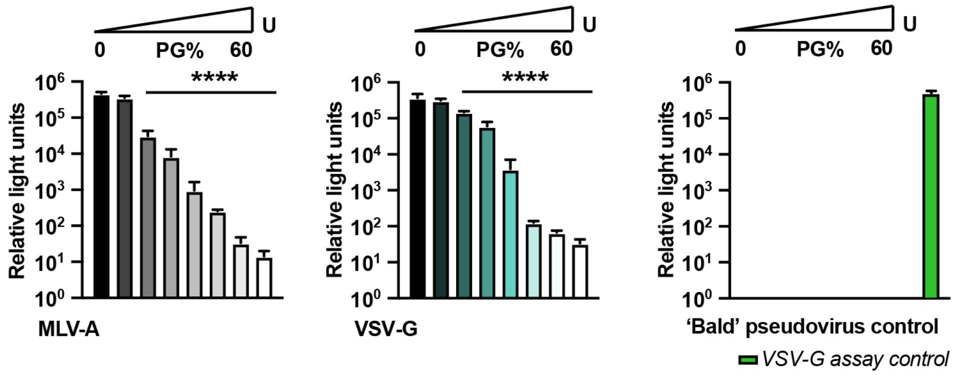


Lentivirus pseudotypes containing different viral glycoproteins from MLV-A (amphotropic murine leukaemia virus) and VSV-G (vesicular stomatis virus), and a 'bald' pseudovirus without viral glycoprotein were incubated with 0-60% [PG] for 5min at RT. Virus infectivity was assessed by firefly luciferase luminescence (N=2, n=3; mean±SD). White bars show background luminescence from mock infected cells. 1-way ANOVA [PG] ****P<0.0001.

**Figure S3. Quantification of PG vapor within the virus transmission tunnel.**

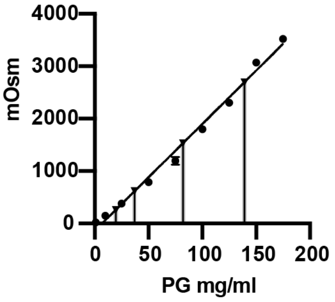


| MicroFogger duration (s) | Av. total [PG] (droplet + vapor) collected in SKC sampler | Av. total [PG] in virus transmission tunnel (mg/L air) |
|---|---|---|
| 5 | 19.57 ± 2.77 | 1.54 ± 0.28 |
| 10 | 37.21 ± 2.20 | 2.93 ± 0.17 |
| 20 | 82.04 ± 9.35 | 6.46 ± 0.74 |
| 30 | 139.09 ± 12.52 | 10.96 ± 0.99 |

PG vapor was introduced into the transmission tunnel using a MicroFogger 2 device activated for different durations (0, 5, 10, 20 and 30s)(N=3 for each duration). Air flow was switched on and total airborne PG within the tunnel (droplets plus vapor) collected in a connected SKC sampler. Osmolarity of the PG samples was assessed (N=3, n=3; mean±SD) and their concentration determined by comparison to the osmolarity of different [PG] standards.

**Figure S4. Virus transmission tunnel: Computational determination of IAV and SARS-CoV-2 plaque area is consistent between ImageJ ColonyArea and ViralPlaque analyses.**

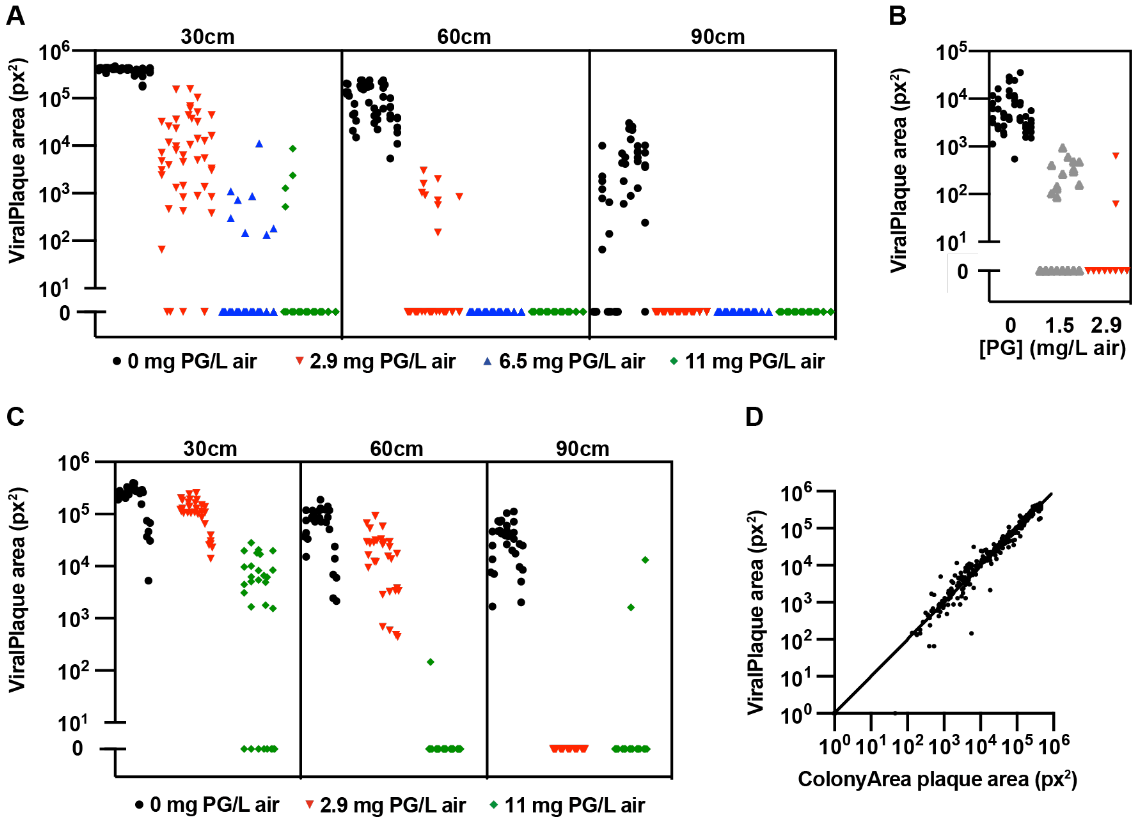


As an *in vitro* model of viral transmission, the IVT has caveats to be considered: nebulised virus droplets are uniform compared to the heterogenous nature of aerosols and respiratory droplets and they also lack respiratory secretions and accompanying salts, glycoproteins and lipids found *in vivo.* (**A)** PG vapor was introduced into the virus transmission tunnel to a final concentration of 0-11 mg/L air prior to nebulization of $10^6$ PFU IAV. Viral plaque area on tissue culture plates at 30, 60 and 90cm from nebulizer was computationally analysed using ImageJ ViralPlaque plugin (N=8, n=6). 2-way ANOVA ([PG] x distance): [PG] ****P<0.0001, distance ****P<0.0001, interaction ****P<0.0001. (**B**). PG vapor was introduced into the virus transmission tunnel to a final concentration of 0-2.9 mg/L air prior to nebulization of $10^4$ PFU IAV. Viral plaque area on plates at 30cm distance from nebulizer was computationally analysed as per (**A**) (N=8, n=6); 1-way ANOVA [PG] ****P<0.0001. (**C**) PG vapor was introduced into the virus transmission tunnel to a final concentration of 0-11 mg PG air prior to nebulization of $3 \times 10^4$ PFU SARS-CoV-2. Viral plaque area was assessed as per (**A**)(N=5, n=6). Differential spacial distribution of IAV and SARS-CoV-2 particles on cell culture plates may reflect specific PG inactivation thresholds or differential infection efficiency of respective permissive cells. 2-way ANOVA [treatment x distance]: treatment ***P<0.001, distance ****P<0.0001, interaction ****P<0.0001. (**D**) Comparison of ColonyArea and ViralPlaque analyses ($R^2$= 0.9648).

**Figure S5. Concomitant nebulization of PG and IAV prevents infection.**

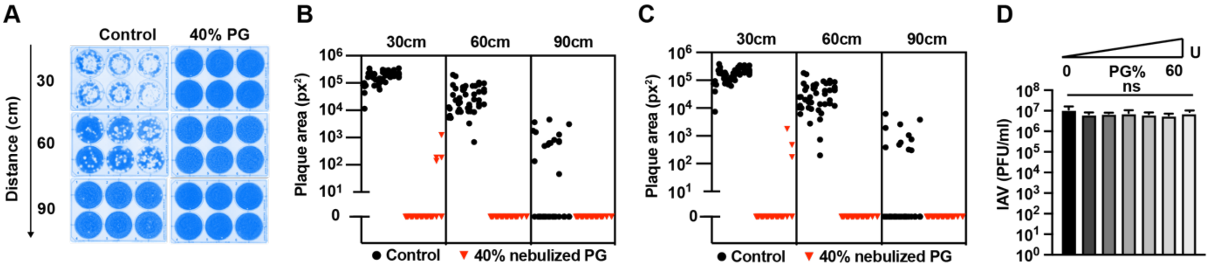


$10^6$ PFU IAV was mixed with PG or PBS control to 40% final concentration and immediately nebulized (<1min incubation) into the virus transmission tunnel. (**A**) Representative plates. Viral plaque area on tissue culture plates at 30cm, 60cm and 90cm were computationally analysed using ImageJ ColonyArea plugin (**B**) and ViralPlaque macros (**C**)(N=8; n=6). 2-way ANOVA ([PG] x distance): [PG] ****P<0.0001, distance ****P<0.0001, interaction ****P<0.0001. (B) (**D**) IAV was mixed with PG to 0-60% final concentration and immediately diluted for plaque assay (N=2; mean±SD)(<1min incubation). 1-way ANOVA [PG] P>0.05.

| Figure | ANOVA | Figure section | | Degree of freedom | F value (DFn, DFd) | Replicates |
|---|---|---|---|---|---|---|
| **1A** | 2-way ANOVA | IAV RT | Interaction | 12 | F (12, 21) = 0.5446 | N= 2 |
| | | | Time | 2 | F (2, 21) = 4.535 | |
| | | | PG concentration | 6 | F (6, 21) = 11.73 | |
| **1B** | 2-way ANOVA | IAV 32 | Interaction | 12 | F (12, 21) = 6.433 | N= 2 |
| | | | Time | 2 | F (2, 21) = 25.07 | |
| | | | PG concentration | 6 | F (6, 21) = 43.59 | |
| **1C** | 2-way ANOVA | IAV 37 | Interaction | 12 | F (12, 21) = 1.087 | N= 2 |
| | | | Time | 2 | F (2, 21) = 0.4154 | |
| | | | PG concentration | 6 | F (6, 21) = 11.67 | |
| **2A** | 2-way ANOVA | SARS-CoV-2 | Interaction | 4 | F (4, 6) = 0.9989 | N= 2 ; n= 4 |
| | | | Time | 2 | F (1.0, 3.0) = 1.002 | |
| | | | PG concentration | 2 | F (2, 3) = 26.72 | |
| **2B** | 2-way ANOVA | EBV | Interaction | 8 | F (8, 15) = 2.118 | N= 2 |
| | | | Time | 4 | F (4, 15) = 1.006 | |
| | | | PG concentration | 2 | F (2, 15) = 24.94 | |
| **2D** | 1-way ANOVA | Ebola pseudovirus | Treatment (between columns) | 7 | F (7, 40) = 151.5 | N=2 ; n= 3 |
| | | | Residual (within columns) | 40 | | |
| | | NL63 coronavirus pseudovirus | Treatment (between columns) | 7 | F (7, 40) = 34.95 | N=2 ; n= 3 |
| | | | Residual (within columns) | 40 | | |
| | | SARS-CoV-1- pseudovirus | Treatment (between columns) | 7 | F (7, 40) = 96.44 | N=2 ; n= 3 |
| | | | Residual (within columns) | 40 | | |
| | | SARS-CoV-2 Wuhan 1 pseudovirus | Treatment (between columns) | 7 | F (7, 40) = 72.37 | N=2 ; n= 3 |
| | | | Residual (within columns) | 40 | | |
| | | SARS-CoV-2 D614G pseudovirus | Treatment (between columns) | 7 | F (7, 40) = 364.2 | N=2 ; n= 3 |
| | | | Residual (within columns) | 40 | | |
| | | SARS-CoV-2 B117 pseudovirus | Treatment (between columns) | 7 | F (7, 40) = 48.13 | N=2 ; n= 3 |
| | | | Residual (within columns) | 40 | | |
| | | VZV- pseudovirus | Treatment (between columns) | 7 | F (7, 40) = 100.1 | N=2 ; n= 3 |
| | | | Residual (within columns) | 40 | | |
| | | Ampho- Pseudovirus | Treatment (between columns) | 7 | F (7, 40) = 381.7 | N=2 ; n= 3 |
| | | | Residual (within columns) | 40 | | |
| | | SARS-CoV-2 Delta pseudovirus | Treatment (between columns) | 7 | F (7, 40) = 261.5 | N=2 ; n= 3 |
| | | | Residual (within columns) | 40 | | |
| | | SARS-CoV-2 Omicron pseudovirus | Treatment (between columns) | 7 | F (7, 40) = 337.3 | N=2 ; n= 3 |
| | | | Residual (within columns) | 40 | | |
| | | MERS pseudovirus | Treatment (between columns) | 7 | F (7, 40) = 641.5 | N=2 ; n= 3 |
| | | | Residual (within columns) | 40 | | |
| | | 229E pseudovirus | Treatment (between columns) | 7 | F (7, 40) = 626.5 | N=2 ; n= 3 |
| | | | Residual (within columns) | 40 | | |
| **3C** | 2-way ANOVA | Plaque Area | Interaction | 6 | F (6, 56) = 167.3 | N= 8 ; n= 6 |
| | | | PG concentration | 3 | F (3, 28) = 367.3 | |
| | | | Distance | 2 | F (1.298, 36.34) =190.9 | |
| **3D** | 1-way ANOVA | Plaque Area | Treatment (between columns) | 2 | F (2, 15) = 43.34 | N= 8 ; n= 6 |
| | | | Residual (within columns) | 15 | | |
| | 1-way ANOVA | Plaque Count | Treatment (between columns) | 2 | F (2, 45) = 157.3 | N= 8 ; n= 6 |
| | | | Residual (within columns) | 45 | | |
| **3F** | 2-way ANOVA | Plaque Area | Interaction | 4 | F (4, 24) = 12.37 | N= 5 ; n= 6 |
| | | | PG concentration | 2 | F (2, 12) = 13.15 | |
| | | | Distance | 2 | F (1.038, 12.45) =53.56 | |
| **3G** | 1-way ANOVA | IAV-Plastic | Treatment (between columns) | 4 | F (4, 13) = 3.903 | N= 4-6 ; n= 3 |
| | | | Residual (within columns) | 13 | | |
| | | IAV-Metal (SS) | Treatment (between columns) | 4 | F (4, 17) = 12.14 | N= 4-6 ; n= 3 |
| | | | Residual (within columns) | 17 | | |
| | | IAV-Metal (A) | Treatment (between columns) | 4 | F (4, 19) = 2.690 | N= 4-6 ; n= 3 |
| | | | Residual (within columns) | | | |
| | | IAV-Glass | Treatment (between columns) | 4 | F (4, 19) = 4.853" | N= 4-6 ; n= 3 |
| | | | Residual (within columns) | | | |
| **S2** | 1-way ANOVA | MLV-A- pseudovirus | Treatment (between columns) | 7 | F (7, 40) = 381.7 | N= 2; n= 3 |
| | | | Residual (within columns) | 40 | | |
| | | VSV-G - pseudovirus | Treatment (between columns) | 7 | F (7, 40) = 100.1 | N= 2; n= 3 |
| | | | Residual (within columns) | 40 | | |
| | | Bald (no glycoproteins) pseudovirus | Treatment (between columns) | 8 | F (8, 45) = 272.4 | N= 2; n= 3 |
| | | | Residual (within columns) | 45 | | |
| **S4A** | 2-way ANOVA | Plaque Area | Interaction | 6 | F (6, 56) = 133.4 | N= 8 ; n= 6 |
| | | | PG concentration | 3 | F (3, 28) = 231.3 | |
| | | | Distance | 2 | F (1.650, 46.21) =155.4 | |
| **S4B** | 1-way ANOVA | Plaque Area | Treatment (between columns) | 2 | F (2, 15) = 43.34 | N= 8 ; n= 6 |
| | | | Residual (within columns) | 15 | | |
| **S4C** | 2-way ANOVA | Plaque Area | Interaction | 4 | F (4, 24) = 10.81 | N= 5 ; n= 6 |
| | | | PG concentration | 2 | F (2, 12) = 12.70 | |
| | | | Distance | 2 | F (1.017, 12.20) =43.32 | |
| **S5A** | 2-way ANOVA | Plaque Area | Interaction | 2 | F (2, 28) = 49.50 | N= 8 ; n= 6 |
| | | | PG concentration | 1 | F (1, 14) = 61.03 | |
| | | | Distance | 2 | F (1.356, 18.99) = 49.54 | |
| **S5C** | 1-way ANOVA | Plaque Count | Treatment (between columns) | 6 | F (6, 7) = 1.04 | N= 2 |
| | | | Residual (within columns) | 7 | | |
| **S5D** | 1-way ANOVA | Plaque Count | Interaction | 2 | F (2, 28) = 38.37 | |
| | | | PG concentration | 1 | F (1, 14) = 44.19 | |
| | | | Distance | 2 | F (1.197, 16.76) = 38.65 | |

**Table S1. Degrees of freedom, replicates and F value for data analysed by ANOVA statistical methods.**
N=biological replicates, n=technical replicates. ANOVA multiple comparisons: Dunnett (1-way) & Tukey (2-way).

### Table S2. Summary of multiple comparison statistical significance.

| Figure | | | | Significance | P value |
|---|---|---|---|---|---|
| 1A | IAV RT | 5 | Control vs. 10% PG | ns | 0.519 |
| | | | Control vs. 20% PG | ns | >0.9999 |
| | | | Control vs. 30% PG | ns | >0.9999 |
| | | | Control vs. 40% PG | ns | 0.8514 |
| | | | Control vs. 50% PG | ns | 0.4061 |
| | | | Control vs. 60% PG | ns | 0.0578 |
| | | 30 | Control vs. 10% PG | ns | 0.946 |
| | | | Control vs. 20% PG | ns | 0.3853 |
| | | | Control vs. 30% PG | ns | 0.3269 |
| | | | Control vs. 40% PG | ns | 0.1275 |
| | | | Control vs. 50% PG | * | 0.0415 |
| | | | Control vs. 60% PG | * | 0.0397 |
| | | 120 | Control vs. 10% PG | ns | >0.9999 |
| | | | Control vs. 20% PG | ns | 0.1667 |
| | | | Control vs. 30% PG | * | 0.0395 |
| | | | Control vs. 40% PG | * | 0.0128 |
| | | | Control vs. 50% PG | ** | 0.008 |
| | | | Control vs. 60% PG | ** | 0.0079 |
| 1B | IAV 32 | 5 | Control vs. 10% PG | ns | 0.5843 |
| | | | Control vs. 20% PG | **** | <0.0001 |
| | | | Control vs. 30% PG | **** | <0.0001 |
| | | | Control vs. 40% PG | **** | <0.0001 |
| | | | Control vs. 50% PG | **** | <0.0001 |
| | | | Control vs. 60% PG | **** | <0.0001 |
| | | 30 | Control vs. 10% PG | ns | 0.9859 |
| | | | Control vs. 20% PG | ns | 0.1021 |
| | | | Control vs. 30% PG | * | 0.0186 |
| | | | Control vs. 40% PG | *** | 0.0002 |
| | | | Control vs. 50% PG | *** | 0.0001 |
| | | | Control vs. 60% PG | *** | 0.0001 |
| | | 120 | Control vs. 10% PG | ns | 0.2709 |
| | | | Control vs. 20% PG | ns | 0.0663 |
| | | | Control vs. 30% PG | * | 0.0209 |
| | | | Control vs. 40% PG | ** | 0.0055 |
| | | | Control vs. 50% PG | ** | 0.0054 |
| | | | Control vs. 60% PG | ** | 0.0054 |
| 1C | IAV 37 | 5 | Control vs. 10% PG | ns | 0.8721 |
| | | | Control vs. 20% PG | ns | >0.9999 |
| | | | Control vs. 30% PG | ns | 0.3684 |
| | | | Control vs. 40% PG | ns | 0.1555 |
| | | | Control vs. 50% PG | ns | 0.1455 |
| | | | Control vs. 60% PG | ns | 0.1431 |
| | | 30 | Control vs. 10% PG | ns | 0.8721 |
| | | | Control vs. 20% PG | * | 0.0367 |
| | | | Control vs. 30% PG | ** | 0.0066 |
| | | | Control vs. 40% PG | ** | 0.0029 |
| | | | Control vs. 50% PG | ** | 0.0029 |
| | | | Control vs. 60% PG | ** | 0.0029 |
| | | 120 | Control vs. 10% PG | * | 0.0123 |
| | | | Control vs. 20% PG | ** | 0.0032 |
| | | | Control vs. 30% PG | *** | 0.0007 |
| | | | Control vs. 40% PG | *** | 0.0006 |
| | | | Control vs. 50% PG | *** | 0.0006 |
| | | | Control vs. 60% PG | *** | 0.0006 |
| 1E | IAV mouse | | Flu Only vs. PG + Flu | ** | 0.0056 |
| 2A | SARS-CoV-2 | 1 | Control vs. 25% PG | ns | 0.6902 |
| | | | Control vs. 50% PG | **** | <0.0001 |
| | | 30 | Control vs. 25% PG | ns | 0.6902 |
| | | | Control vs. 50% PG | **** | <0.0001 |
| 2B | EBV | 5 | Control vs. 25% PG | ns | 0.9259 |
| | | | Control vs. 50% PG | ns | 0.1517 |
| | | 15 | Control vs. 25% PG | ** | 0.0065 |
| | | | Control vs. 50% PG | ** | 0.0058 |
| | | 30 | Control vs. 25% PG | * | 0.0136 |
| | | | Control vs. 50% PG | * | 0.0136 |
| 2D | 229E- coronavirus pseudovirus | | 0% PG vs. 10% PG | **** | <0.0001 |
| | | | 0% PG vs. 20% PG | **** | <0.0001 |
| | | | 0% PG vs. 30% PG | **** | <0.0001 |
| | | | 0% PG vs. 40% PG | **** | <0.0001 |
| | | | 0% PG vs. 50% PG | **** | <0.0001 |
| | | | 0% PG vs. 60% PG | **** | <0.0001 |
| | NL63-coronavirus pseudovirus | | 0% PG vs. 10% PG | ns | 0.9994 |
| | | | 0% PG vs. 20% PG | **** | <0.0001 |

| | | | | | |
|---|---|---|---|---|---|
| | | | 0% PG vs. 30% PG | **** | <0.0001 |
| | | | 0% PG vs. 40% PG | **** | <0.0001 |
| | | | 0% PG vs. 50% PG | **** | <0.0001 |
| | | | 0% PG vs. 60% PG | **** | <0.0001 |
| | SAR1 coronavirus | | 0% PG vs. 10% PG | **** | <0.0001 |
| | | | 0% PG vs. 20% PG | **** | <0.0001 |
| | | | 0% PG vs. 30% PG | **** | <0.0001 |
| | | | 0% PG vs. 40% PG | **** | <0.0001 |
| | | | 0% PG vs. 50% PG | **** | <0.0001 |
| | | | 0% PG vs. 60% PG | **** | <0.0001 |
| | MERS coronavirus | | 0% PG vs. 10% PG | * | 0.0195 |
| | | | 0% PG vs. 20% PG | **** | <0.0001 |
| | | | 0% PG vs. 30% PG | **** | <0.0001 |
| | | | 0% PG vs. 40% PG | **** | <0.0001 |
| | | | 0% PG vs. 50% PG | **** | <0.0001 |
| | | | 0% PG vs. 60% PG | **** | <0.0001 |
| | Ebola pseudovirus | | 0% PG vs. 10% PG | **** | <0.0001 |
| | | | 0% PG vs. 20% PG | **** | <0.0001 |
| | | | 0% PG vs. 30% PG | **** | <0.0001 |
| | | | 0% PG vs. 40% PG | **** | <0.0001 |
| | | | 0% PG vs. 50% PG | **** | <0.0001 |
| | | | 0% PG vs. 60% PG | **** | <0.0001 |
| | SARS-CoV-2 Wuhan 1 | | 0% PG vs. 10% PG | ns | >0.9999 |
| | | | 0% PG vs. 20% PG | **** | <0.0001 |
| | | | 0% PG vs. 30% PG | **** | <0.0001 |
| | | | 0% PG vs. 40% PG | **** | <0.0001 |
| | | | 0% PG vs. 50% PG | **** | <0.0001 |
| | | | 0% PG vs. 60% PG | **** | <0.0001 |
| | SARS-CoV-2 D614G | | 0% PG vs. 10% PG | * | 0.0396 |
| | | | 0% PG vs. 20% PG | **** | <0.0001 |
| | | | 0% PG vs. 30% PG | **** | <0.0001 |
| | | | 0% PG vs. 40% PG | **** | <0.0001 |
| | | | 0% PG vs. 50% PG | **** | <0.0001 |
| | | | 0% PG vs. 60% PG | **** | <0.0001 |
| | SARS-CoV-2 Alpha | | 0% PG vs. 10% PG | * | 0.0423 |
| | | | 0% PG vs. 20% PG | **** | <0.0001 |
| | | | 0% PG vs. 30% PG | **** | <0.0001 |
| | | | 0% PG vs. 40% PG | **** | <0.0001 |
| | | | 0% PG vs. 50% PG | **** | <0.0001 |
| | | | 0% PG vs. 60% PG | **** | <0.0001 |
| | SARS-CoV-2 Delta | | 0% PG vs. 10% PG | ns | 0.2651 |
| | | | 0% PG vs. 20% PG | **** | <0.0001 |
| | | | 0% PG vs. 30% PG | **** | <0.0001 |
| | | | 0% PG vs. 40% PG | **** | <0.0001 |
| | | | 0% PG vs. 50% PG | **** | <0.0001 |
| | | | 0% PG vs. 60% PG | **** | <0.0001 |
| | SARS-CoV-2 Omicron | | 0% PG vs. 10% PG | ns | 0.9309 |
| | | | 0% PG vs. 20% PG | **** | <0.0001 |
| | | | 0% PG vs. 30% PG | **** | <0.0001 |
| | | | 0% PG vs. 40% PG | **** | <0.0001 |
| | | | 0% PG vs. 50% PG | **** | <0.0001 |
| | | | 0% PG vs. 60% PG | **** | <0.0001 |
| 3C | | Plate 1 | 0 vs. 2.93 PG mg/L air | **** | <0.0001 |
| | | | 0 vs. 6.46 PG mg/L air | **** | <0.0001 |
| | | | 0 vs. 10.593 PG mg/L air | **** | <0.0001 |
| | | Plate 2 | 0 vs. 2.93 PG mg/L air | **** | <0.0001 |
| | | | 0 vs. 6.46 PG mg/L air | **** | <0.0001 |
| | | | 0 vs. 10.593 PG mg/L air | **** | <0.0001 |
| | | Plate 3 | 0 vs. 2.93 PG mg/L air | ns | 0.9843 |
| | | | 0 vs. 6.46 PG mg/L air | ns | 0.9843 |
| | | | 0 vs. 10.593 PG mg/L air | ns | 0.9843 |
| 3D | Plaque area | Plate 1 | 0 vs. 1.541 PG mg/L air | **** | <0.0001 |
| | | | 0 vs. 2.93 PG mg/L air | **** | <0.0001 |
| | Plaque count | Plate 1 | 0 vs. 1.541 PG mg/L air | **** | <0.0001 |
| | | | 0 vs. 2.93 PG mg/L air | **** | <0.0001 |
| 3F | | Plate 1 | 0 vs. 2.93 PG mg/L air | ** | 0.0061 |
| | | | 0 vs. 10.593 PG mg/L air | **** | <0.0001 |
| | | Plate 2 | 0 vs. 2.93 PG mg/L air | ns | 0.0793 |
| | | | 0 vs. 10.593 PG mg/L air | * | 0.0105 |
| | | Plate 3 | 0 vs. 2.93 PG mg/L air | ns | 0.3127 |
| | | | 0 vs. 10.593 PG mg/L air | ns | 0.3182 |
| 3G | | Plastic | 0 vs. 2.93 PG mg/L air | ** | 0.0027 |

| | | | | | |
|---|---|---|---|---|---|
| | | | 0 vs. 6.46 PG mg/L air | *** | 0.0002 |
| | | | 0 vs. 10.593 PG mg/L air | **** | <0.0001 |
| | | Stainless steel | 0 vs. 2.93 PG mg/L air | ns | 0.0554 |
| | | | 0 vs. 6.46 PG mg/L air | * | 0.0434 |
| | | | 0 vs. 10.593 PG mg/L air | * | 0.0239 |
| | | Glass | 0 vs. 2.93 PG mg/L air | * | 0.0434 |
| | | | 0 vs. 6.46 PG mg/L air | ** | 0.0054 |
| | | | 0 vs. 10.593 PG mg/L air | ** | 0.0053 |
| | | Aluminium | 0 vs. 2.93 PG mg/L air | ns | 0.1401 |
| | | | 0 vs. 6.46 PG mg/L air | * | 0.0415 |
| | | | 0 vs. 10.593 PG mg/L air | * | 0.0402 |
| S2 | MLV-A pseudovirus | | 0% PG vs. 10% PG | **** | <0.0001 |
| | | | 0% PG vs. 20% PG | **** | <0.0001 |
| | | | 0% PG vs. 30% PG | **** | <0.0001 |
| | | | 0% PG vs. 40% PG | **** | <0.0001 |
| | | | 0% PG vs. 50% PG | **** | <0.0001 |
| | | | 0% PG vs. 60% PG | **** | <0.0001 |
| | VSV-G pseudovirus | | 0% PG vs. 10% PG | ns | 0.0559 |
| | | | 0% PG vs. 20% PG | **** | <0.0001 |
| | | | 0% PG vs. 30% PG | **** | <0.0001 |
| | | | 0% PG vs. 40% PG | **** | <0.0001 |
| | | | 0% PG vs. 50% PG | **** | <0.0001 |
| | | | 0% PG vs. 60% PG | **** | <0.0001 |
| | 'Bald' pseudovirus control | | 0% PG vs. 10% PG | ns | >0.9999 |
| | | | 0% PG vs. 20% PG | ns | >0.9999 |
| | | | 0% PG vs. 30% PG | ns | >0.9999 |
| | | | 0% PG vs. 40% PG | ns | >0.9999 |
| | | | 0% PG vs. 50% PG | ns | >0.9999 |
| | | | 0% PG vs. 60% PG | ns | >0.9999 |
| S4A | | Plate 1 | 0 vs. 2.93 PG mg/L air | **** | <0.0001 |
| | | | 0 vs. 6.46 PG mg/L air | **** | <0.0001 |
| | | | 0 vs. 10.593 PG mg/L air | **** | <0.0001 |
| | | Plate 2 | 0 vs. 2.93 PG mg/L air | * | 0.018 |
| | | | 0 vs. 6.46 PG mg/L air | * | 0.0178 |
| | | | 0 vs. 10.593 PG mg/L air | * | 0.0178 |
| | | Plate 3 | 0 vs. 2.93 PG mg/L air | ns | 0.2653 |
| | | | 0 vs. 6.46 PG mg/L air | ns | 0.2653 |
| | | | 0 vs. 10.593 PG mg/L air | ns | 0.2653 |
| S4B | Plaque area | Plate 1 | 0 vs. 1.541 PG mg/L air | **** | <0.0001 |
| | | | 0 vs. 2.93 PG mg/L air | **** | <0.0001 |
| S4C | | Plate 1 | 0 vs. 2.93 PG mg/L air | ns | 0.2119 |
| | | | 0 vs. 10.593 PG mg/L air | * | 0.0229 |
| | | Plate 2 | 0 vs. 2.93 PG mg/L air | ns | 0.112 |
| | | | 0 vs. 10.593 PG mg/L air | * | 0.0396 |
| | | Plate 3 | 0 vs. 2.93 PG mg/L air | * | 0.0367 |
| | | | 0 vs. 10.593 PG mg/L air | * | 0.0381 |
| S5A | | Plate 1 | 0% vs 40% | **** | <0.0001 |
| S5B | | Plate 1 | 0% vs 40% | **** | <0.0001 |
| S5C | IAV | | 0% PG vs. 10% PG | ns | 0.3435 |
| | | | 0% PG vs. 20% PG | ns | 0.4395 |
| | | | 0% PG vs. 30% PG | ns | 0.5515 |
| | | | 0% PG vs. 40% PG | ns | 0.3435 |
| | | | 0% PG vs. 50% PG | ns | 0.2029 |
| | | | 0% PG vs. 60% PG | ns | 0.5515 |
